# Designer micro/nano-crumpled MXene multilayer coatings accelerate osteogenesis and regulate macrophage polarization

**DOI:** 10.1101/2024.01.10.574996

**Authors:** Mohammad Asadi Tokmedash, Jouha Min

**Affiliations:** Department of Chemical Engineering, University of Michigan, Ann Arbor, MI 48109; Department of Biomedical Engineering, University of Michigan, Ann Arbor, MI 48109; Department of Macromolecular Science and Engineering, University of Michigan, Ann Arbor, MI 48109; Weil Institute for Critical Care Research and Innovation, University of Michigan, Ann Arbor, MI 48109

**Keywords:** Micro-/nanoscale surface topography, crumpled structures, MXene, Layer-by-Layer, osteogenesis, immunomodulation, tissue engineering

## Abstract

Effective tissue regeneration and immune responses are essential for the success of biomaterial implantation. Although the interaction between synthetic materials and biological systems is well-recognized, the role of surface topographical cues in regulating the local osteoimmune microenvironment—specifically, their impact on host tissue and immune cells and their dynamic interactions—remains underexplored. This study addresses this gap by investigating the impact of surface topography on osteogenesis and immunomodulation. We fabricated MXene/Hydroxyapatite (HAP)-coated surfaces with controlled 2.5D nano-, submicro-, and micro-scale topographical patterns using our custom bottom-up pattering method. These engineered surfaces were employed to assess the behavior of osteoblast precursor cells and macrophage polarization. Our results demonstrate that MXene/HAP-coated surfaces with microscale crumpled topography significantly influence osteogenic activity and macrophage polarization: These surfaces notably enhanced osteoblast precursor cell spreading, proliferation, differentiation, and facilitated a shift in macrophages towards an anti-inflammatory, pro-healing M2 phenotype. The observed cell responses indicate that the physical cues from the crumpled topographies, combined with the chemical cues from the MXene/HAP coatings, synergistically create a favorable osteoimmune microenvironment. This study presents the first evidence of employing MXene/HAP-multilayer coated surfaces with finely crumpled topography to concurrently facilitate osteogenesis and immunomodulation for improved implant-to-tissue integration. The tunable topographic patterns of these coatings, coupled with a facile and scalable fabrication process, make them widely applicable for various biomedical purposes. Our results highlight the potential of these novel coatings to improve the *in vivo* performance and fate of implants by modulating the host response at the material interface.

## INTRODUCTION

The growing global aging population and rising incidence of tissue-degenerative diseases have led to a dramatic increase in the demand for implantable biomaterials and devices over the past few decades.^1^ Despite the growing use of implants, we still face key challenges, including adverse immunological foreign-body responses (FBR) and inadequate tissue integration. Implant failure rates due to FBR are conservatively estimated to be >10% (and 30% for breast implants), while aseptic loosening accounts for ∼55% of hip and 31% of knee joint arthroplasty failures.^2-6^ To address these issues, recent studies have focused on harnessing the plasticity and adaptive nature of stem cells and innate immune cells.^7^ These cells can dynamically alter their phenotypes in response to local environmental cues. In orthopedic implants, for example, the proliferation and differentiation of osteoblast precursor cells is vital for bone formation and implant stability. This biological process is closely linked with the establishment of a favorable osteoimmune microenvironment, which can be facilitated by the polarization of macrophages—a key player of the innate immune system—towards an anti-inflammatory and prohealing M2 phenotype.^8-10^ Both tissue regeneration and immunomodulation are integral to the overall success of implant-to-bone integration, highlighting the importance of understanding of these biological interactions at the material interfaces.

In many biomedical applications, the interaction between synthetic materials and biological systems is crucial. Biophysical and biochemical factors of these material surfaces act as potent regulators of cell activity. In particular, surface topography provides a profound physical cue to modulate their interaction with biological entities. For example, microtopography, such as microgratings, guide cell shape and motility, while nanotopography affects anchorage-dependent cells by targeting transmembrane receptor proteins involved in intracellular signal transduction.^11, 12^ Remarkably, cells can sense and interact with substrate features as small as 5 nm.^13^ Recent research indicates that surfaces with topographical patterns, as opposed to smooth surfaces, enhance cell adhesion, proliferation, and differentiation, and are effective in regulating macrophage polarization.^14, 15^ However, despite these advancements, there remains a significant knowledge gap in the field. As outlined in **Table S1**, most research has focused on either tissue/stem cells or immune cells, overlooking the three-body interactions that are fundamental to the pathophysiology to implant-to-tissue osteointegration. Furthermore, the current topographic patterning methods such as lithographic patterning of nanogratings are complicated, expensive, and not readily scalable, which hamper the translation of the research findings into practice.

To address these research gaps, our study aimed to elucidate the responses of preosteoblast cells and macrophages to controlled surface topographical cues (**Scheme 1**). We implemented a novel bottom-up topographic patterning method to create surfaces with precisely tuned nano- to micro-scale peaks and valleys. This patterning method involved an emerging mechanical nanomanufacturing, specifically by shrinking a stiff coating on a compliant substrate and generating 2.5D crumpled structures.^16, 17^ Compared to the conventional lithographic methods, this method has advantages in the simplicity and scalability of fabrication. A key aspect of our approach was the deposition of a 2D nanomaterial (MXene) and an inorganic nanoparticle (Hydroxyapatite, HAP). MXene, a group of transition metal carbides/nitrides/carbonitrides, has been recognized for its high biocompatibility and osteoconductive properties, presenting promising potential in biomedical applications.^18, 19^ HAP (Ca_5_(PO_4_)_3_(OH)), a key component of human hard tissue, is renowned for its high osteoconductivity, making it an ideal material candidate for orthopedic uses.^20^ The atomically thin properties of MXene allowed for the creation of ultra-thin films using the layer-by-layer (LbL) fabrication method. This method offers several advantages, including precise nanoscale control, straightforwardness in application, and the potential for scalability, aligning perfectly with our research objectives.^21^

We hypothesized that MXene/HAP multilayer coatings with controlled topographies would modulate mechanotransduction signaling pathways—such as RhoA/Rho–associated protein kinase signaling—thereby regulating the activity of adhered cells. In this study, we aim to elucidate the impact of controlled surface topography on osteoblast precursor cells and macrophages, ultimately establishing an optimal osteoimmune microenvironment for implant-to-bone integration. Our approach integrated the use of osteoconductive materials (chemical cues) and precisely engineered topographies (physical cues). To our knowledge, this study is first to demonstrate the application of finely crumpled (MXene/HAP)_*n*_ multilayer coatings in facilitating both osteogenesis and immunomodulation—particularly, macrophage polarization—to enhance implant-to-tissue integration. In our methodology, we first fabricated a conformal (MXene/HAP)_*n*_ multilayer coating on a biaxially-oriented polystyrene (PS) substrate using LbL assembly. Subsequent thermal annealing induced the mechanical deformation of the PS film, resulting in a distinctive “crumpled” structure within the (MXene/HAP)_*n*_ multilayer coating. The morphological characteristics of these crumpled MXene/HAP coatings were analyzed using scanning electron microscopy (SEM) and atomic force microscopy (AFM). Following this, we conducted a comprehensive examination of the behavior of osteoblast precursor cells on diverse MXene/HAP-coated surfaces with controlled topographies. Additionally, we explored the interactions between these engineered surfaces and macrophages. For comparative analysis, we included both uncoated and MXene/HAP-coated planar surfaces as controls to evaluate the distinct impacts of topographical cues and chemical cues on cell responses. Our findings indicate that MXene/HAP-coated surfaces with microscale crumpled topography significantly influence osteogenic activity and macrophage polarization, potentially contributing to the formation of a favorable osteoimmune microenvironment. Notably, the nano-/micro-structure of these surfaces are highly tunable, and the fabrication process is scalable for different materials and substrates.

**Scheme 1.**
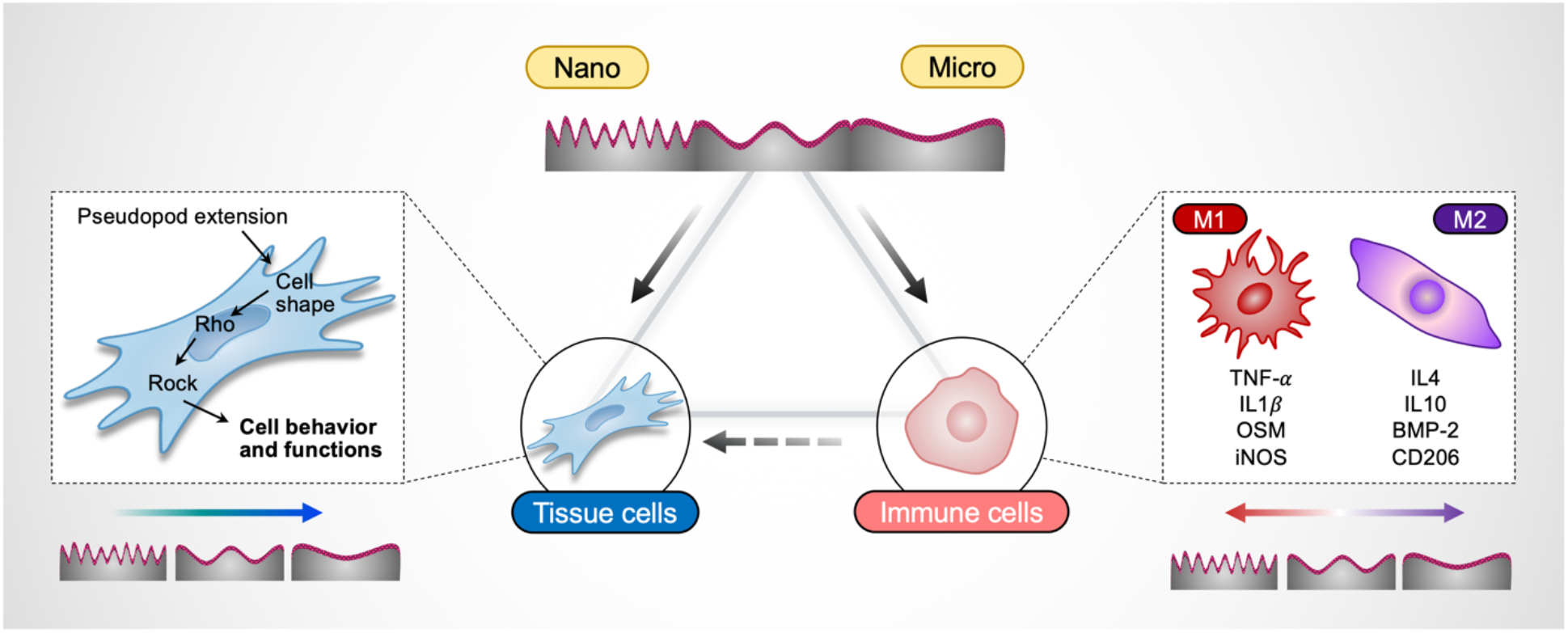
This study focuses on the three-body interactions between topographically patterned materials, tissue cells, and immune cells. We hypothesize that surface topographical cues can facilitate osteointegration by properly regulating the osteoimmune tissue.

## RESULTS

### System design and fabrication

**Fig. 1a** illustrates the bottom-up fabrication process for our osteophilic MXene multilayers coatings with crumpled topographical patterns. We fabricated multilayer films by sequential LbL deposition of Ti_3_CT_x_ MXene nanosheets, produced via the selective etching of aluminum atoms from Ti_3_Al_2_C_2_ MAX phase using a mixture of HCl and LiF solution (**Fig. S1**)^22^, and hydroxyapatite (HAP) nanoparticles on a polystyrene (PS) substrate, leveraging hydrogen bonding for self-assembly. The hydrogen bonding occurs between active hydroxyl groups (-OH) in HAP and functional groups (-F, =O, -OH) in MXene nanosheets, promoting self-assembly processes (**Fig. S2**).^23^ The multilayer films are denoted as (MXene/HAP)_*n*_, where *n* corresponds to the number of bilayers that are repeated in the LbL deposition process. After the fabrication of the MXene multilayer film on the PS substrate, we applied quick thermal annealing. This process causes the PS substrate to shrink and lead to the formation of a controlled topographical pattern characterized by nano to micro-scale peaks and valleys.

**Figure 1.**
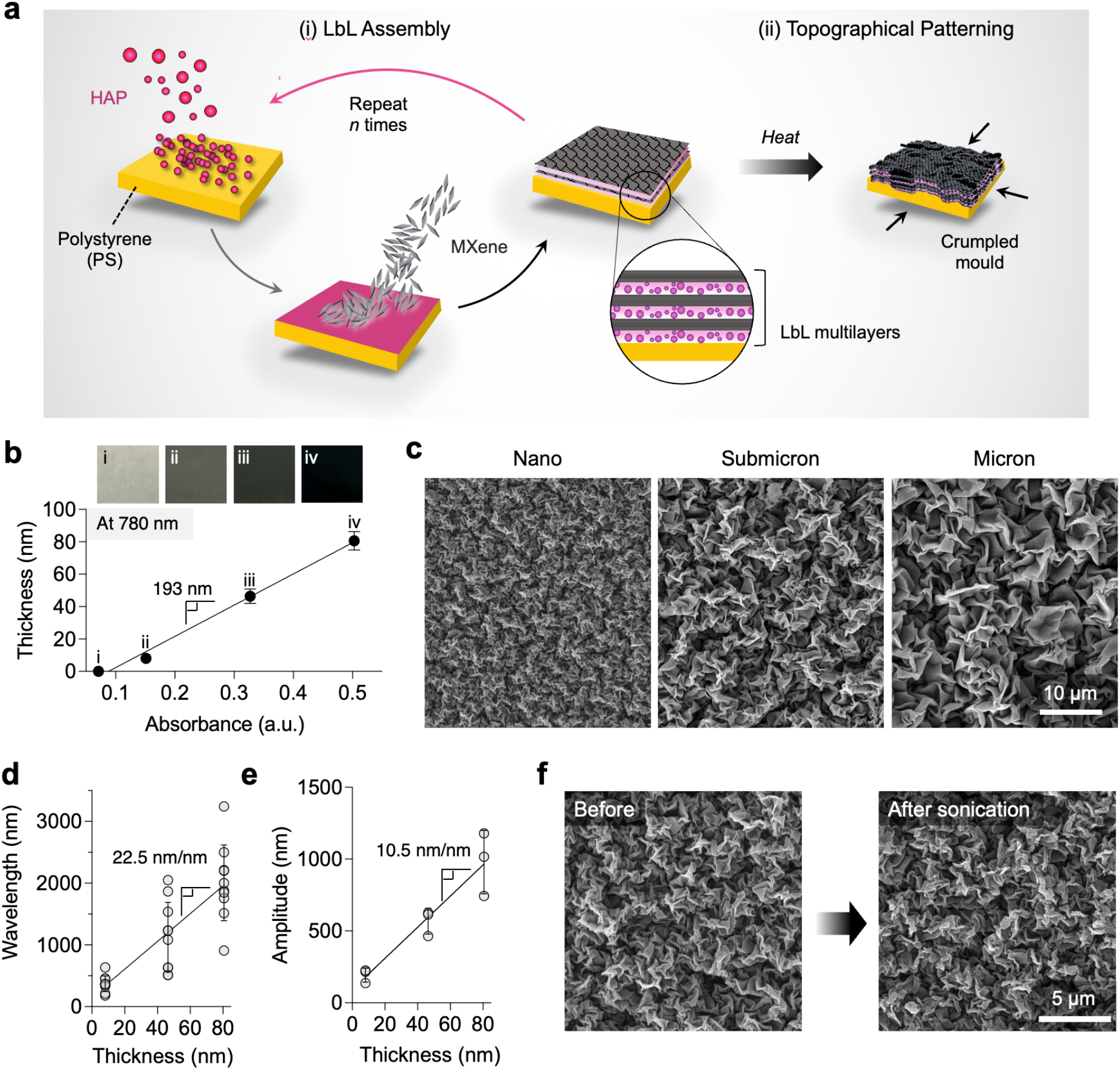
Structural characterization of as-prepared (MXene/HAP)_*n*_ multilayers. **(a)** Schematic of (i) LbL nanofabrication of (MXene/HAP)_*n*_ multilayers on biaxially-oriented PS films and (ii) crumpled topography formation. **(b)** Growth curve of the films as a function of absorbance and digital photos of (i) uncoated PS films, and (ii) 1 bilayer (iii) 10 bilayers, and (iv) 20 bilayers of (MXene/HAP)_*n*_ multilayers. **(c)** SEM images of the crumpled (MXene/HAP) multilayers. **(d)** Wavelength and **(e)** peak-to-peak amplitude of the crumpled (MXene/HAP) multilayers as a function of film thickness. **(f)** Testing of mechanical stability (longevity) of the microstructure through a 15-minute batch sonication in water. Each condition was tested with at least three replicates (error bar = standard deviations; *n* ≥ 3).

We chose MXene nanosheets as the building block for our LbL and subsequent nanomanufacturing processes because of their high aspect-ratio with atomic thickness, high charge density, high water solubility, and high mechanical properties.^24^ These characteristics make them particularly suitable for our engineering and biological objectives. Recently, MXene has been reported to have osteoconductive properties.^18, 19, 25^ We also selected Hydroxyapatite (HAP, Ca_10_(PO_4_)_6_(OH)_2_)-the principal inorganic component of human hard tissues such as cortical bone and teeth to which it has a high bonding affinity.^26^ In the LbL assembly, it serves as a complementary species of MXene, imparting osteoconductive properties (i.e., capability to specifically support the attachment and proliferation of osteoprogenitor cells) to the coatings.^27^ Despite its wide use in hard tissue regeneration, the relatively low mechanical strength of standard HAP ceramics limits its application to low load-bearing areas. Recently, HAP nanoparticles are considered ideal biomaterials thanks to their excellent biocompatibility and bone integration ability.^28^ Numerous studies have shown their role in enhancing stem cell proliferation and differentiation in bone defects, highlighting their potential for regenerative medicine.^29-31^

To explore the growth behavior of our osteophilic multilayer films, we used an immersive LbL self-assembly technique to develop (MXene/HAP)_*n*_ multilayers on PS sheets and glass. As the number of bilayer cycles increased, the coating, deriving its color from the MXene sheets, successfully darkened (**Fig. 1b**). We also measured the thickness of (MXene/HAP)_*n*_ multilayers from the AFM images of the films grown on glass. The film thickness exhibits a linear increase versus adsorption at 780 nm, consistent with the MXene nanosheet (**Fig. 1b**). The linear growth trend is a feature of the successful LbL self-assembly process, indicating complete alternation of the two components during the layering procedure. Notably, we observed a direct relationship exists between film thickness and the size of HAP nanoparticles; larger HAP nanoparticles tend to accelerate film growth (**Fig. S3**).

### Controlled topographical patterning and characterization

After fabricating an MXene multilayer film on a PS substrate, we applied a quick thermal annealing at 130°C, above the glass transition temperature of PS (∼100°C), to cause the substrate to shrink by releasing the prestretched strain. Due to the modulus mismatch between the MXene multilayers coating and the PS substrate, this thermal shrinkage led to isotropic crumpling of the MXene LbL films. The strain defined as ε = (*L*_i_ – *L*_f_)/*L*_i_, where *L*_i_ and *L*_f_ are the lengths of the PS films before and after the shrinkage process, respectively. This crumpling can be further transformed into ridge-like structures at high shrinkage rates.^32, 33^

We examined the morphological details of the crumpled MXene LbL films using SEM and AFM. SEM images show diverse crumpled structures at varying length scales and densities (**Fig. 1c**). Typically, these crumpled structures are characterized by wavelength (*λ*, distance between peak to peak of periodic ripples) and amplitude (*A*, height difference between peaks).^34^ By controlling the thickness of the MXene multilayer coating at a high deformation strain (ε = 0.9), these characteristic parameters can be finely tuned. As per classical buckling theory, a thin stiff coating with thickness *h* deposited on a uniaxially prestretched substrate will buckle with a characteristic wavelength of *λ* = 2π*h*(*Ē*_c_/3*Ē*_s_)^1/3^ where the plane-strain elastic modulus *Ē*_*i*_ = *E*_*i*_/(1 − *ν*_i_^2^) is given in terms of the Young’s modulus *E* and Poisson’s ratio *v*_i_ of the multilayer coating (c) or PS substrate(s), respectively.^35^ This theory gives qualitative insight into the isotropic crumpled structures formed here through biaxial deformation. In this study, we analyzed SEM and AFM data to estimate the characteristic features of the crumpled (MXene/HAP)_*n*_ multilayers with different thicknesses (**Fig. 1c** and **Fig. S4**). Note that for amplitude, we assumed the ridge-like structure of the MXene LbL film has a triangle waveform and used the relationship: 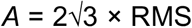 (root-mean-square). Our results show a nearly increase in feature sizes as film thickness increases from 8.07 nm to 80.6 nm: (i) *λ* from 372 nm to 2010 nm (**Fig. 1c-d**) and (ii) *A* from 193 nm to 979 nm (**Fig. 1e** and **Fig. S3**). The fabrication of these diverse nanostructured surfaces with finely crumpled configurations proved to be scalable (i.e. capable to scale up in size) and robust. For subsequent biological studies, we categorized the crumpled multilayers into three distinct groups based on their characteristic wavelengths: Nano (200-400 nm), Submicro (900-1100 nm), and Micro (1900-2100 nm). We specifically designed the topographical cues on a subcellular scale to enable focal adhesions and cell protrusions to initiate dynamic, actin-based subcellular processes.^36^

To ensure the long-term functionality of the engineered surfaces, we evaluated the mechanical stability of crumpled MXene/HAP coatings on PS substrates by characterizing interfacial failure under ultrasonication. After 15 minutes of ultrasonication in water, the crumpled MXene/HAP coatings demonstrated robust stability (i.e., negligible morphological distortion) (**Fig. 1f** and **Fig. S5**). Moreover, the crumpled MXene/HAP coatings maintained their integrity without degradation, even after 30 days of incubation in cell culture media (**Fig. S6**).

### Morphology, spreading and proliferation of preosteoblast cells

Enhancing cell adhesion and proliferation on the implant’s surfaces is crucial for the repair of damaged bone tissue at the implant site.^37^ Successful osteointegration depends on the migration and attachment of osteoprogenitor cells to the damaged area, guided by chemoattractive signals.^38^ Accordingly, we hypothesized that incorporating HAP nanoparticles into the system would form a favorable osteoconductive microenvironment.^20^ Additionally, the surface topographical cues would provide essential mechanotransduction signals to facilitate the proliferation and differentiation of cells into mineral depositing osteoblasts.^39, 40^ Based on the literature, we anticipated that microscale crumpled topography would more significantly enhance cell proliferation and differentiation compared to their nanoscale counterparts.^41-43^ To test this hypothesis, we assessed cell morphology, proliferation, and differentiation of murine preosteoblast MC3T3-E1 cells on various samples. The test samples included MXene multilayers featuring crumpled topographical patterns ranging from nano to micro scale: namely Nano, Submicro, and Micro groups. For controls, we used both uncoated and MXene/HAP-coated planar PS substrates to evaluate the impact of topographical cues and chemical cues on cellular responses (**Fig. 2a**).

**Figure 2.**
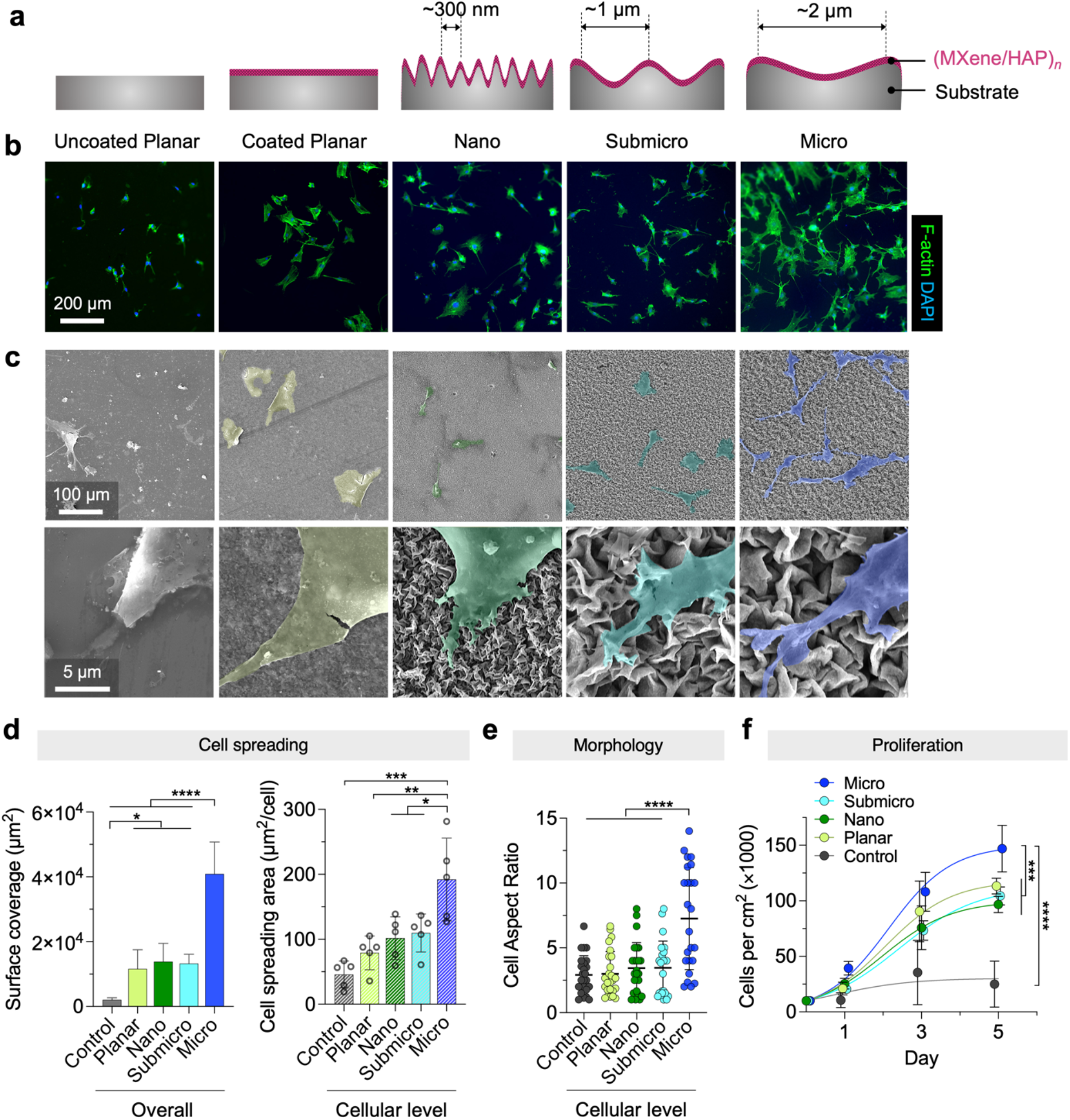
Cell adhesion, spreading, and proliferation of osteoblast precursor cells on flat controls and crumpled (MXene/HAP) multilayer films. **(a)** Schematic illustration of test samples with different coatings and topographic parameters (wavelength, amplitude). **(b)** Merged fluorescent images of cell spreading morphologies after culturing for 24 hours, F-actin (green) and nucleus (blue). **(c)** SEM images of cells on different samples after culturing for 24 hours. **(d)** Overall surface coverage and individual cell spreading area analyzed on fluorescent images using Cellprofiler. **(e)** Relative cell aspect ratio analyzed on SEM images. **(f)** Cell proliferation rate of MC3T3-E1 on different samples for 1, 3, and 5 days. Each condition was tested with at least three biological replicates (error bar = standard deviations; *n* ≥ 5; \*\*\**\*p* < 0.0001, \**\**\**p* < 0.001, *and* \**p* < 0.*05*), and at least three random images were taken from each sample.

First, we investigated cell morphology and spreading of preosteoblast cells on different groups after 24 hours of culturing, using immunofluorescence (IF) imaging and SEM. IF analysis involved staining cytoskeleton actin (green), and nucleus (blue) of preosteoblast cells. Our results show robust cell adhesion on all samples, albeit with varied spreading across different surface topographies (**Fig. 2b-c**). Overall, cells cultured on the MXene/HAP-coated surfaces with topographical patterns displayed significantly enhanced surface coverage compared to the uncoated planar control (**Fig. 2d**). Cells on the coated planar surfaces were observed to adopt a more prominent round-to-polygonal shape with thick and flat lamellipodia and smooth boundaries. This group noted a 5.6-fold increase in cell surface coverage compared to the uncoated planar control, indicating excellent biocompatibility and osteoconductive properties of the MXene/HAP multilayer coatings (**Fig. 2d**). In the Nano and Submicro groups, cells exhibited slender and elongated shape with enhanced filopodia formation, showing a >6.4-fold increase in cell surface coverage compared to the uncoated planar control. Cells in the Micro group exhibit a mature osteoblast-like appearance, characterized by a stellate shape^44, 45^ and elongated cytoplasmic processes with lamellipodia extending and cellular pseudopodia beyond the cell body and numerous filopodia anchoring to the surface protrusions. This collective cellular behavior resulted in a significant increase in surface coverage, 20-fold greater than the uncoated planar control (40900 ± 9860 μm^2^ for the Micro vs. 2060 ± 653 μm^2^ for the uncoated planar control, *p* < 0.0001) and 3-fold greater than the Nano and Submicro groups.

We also analyzed cell spreading and morphology at a single cell level. Using CellProfilier software, we processed the individual cell spreading area on fluorescent staining of F-actin and nucleus post 24-hour culturing. We further analyze cell aspect ratio, using SEM images. Our results reveal increased cell spreading area and aspect ratio for crumpled surfaces with microscale topography compared to other groups (**Fig. 2d-e**). Specifically, the extent of spreading for individual cells in the Micro group was nearly double compared to the Nano and Submicro structures, and approximately four times that of the uncoated planar control. Similarly, the cell aspect ratio for cells in the Micro group was found to be twice as large compared to those in the Nano and Submicro structures, and nearly 2.5 times larger than those in the uncoated planar control.

Next, we assessed the proliferation of preosteoblast cells on different surfaces using IF analysis. As shown **Fig. 2f**, all coated samples demonstrated significant cell proliferation from day 1 to day 5, regardless of their topographical patterns, compared to the uncoated planar control. Notably, on day 5, the Micro group showed a substantially higher proliferation rate than other groups: with counts of 147 ± 21.0 (×10^3^) cells for the Micro vs. 96.8 ± 7.32 (×10^3^) cells for the Nano (*p* = 0.0002), 113 ± 7.12 (×10^3^) cells for the coated planar control (*p* = 0.0078), and 24.9 ± 20.8 (×10^3^) cells for the uncoated planar control (*p* < 0.0001). The significant increases in cell proliferation as well as adhesion (i.e., surface coverage and spreading) within the Micro group support our hypothesis that microscale crumpled topography more notably enhances cell proliferation compared to their nanoscale counterparts. In summary, the initial behavior and responses of preosteoblast cells showed noticeable differences between the planar and patterned groups, as well as between the uncoated and coated groups. Such differences could potentially impact the osteogenic differentiation of the preosteoblast cells.

### Osteogenic differentiation

We furthered our investigations to explore how MXene/HAP-coated surfaces with crumpled topographical patterns influence the osteogenic differentiation. To this end, we tested alkaline phosphatase (ALP) activity—an early marker of induction of bone differentiation—in MC3T3-E1 cells (**Fig. 3a**). Briefly, Cells were cultured on different samples and exposed to osteogenic induction media (growth medium supplemented with L-ascorbic acid, -glycerol phosphate and dexamethasone). At 7 and 14 days, for ALP assessment through a colorimetric assay, the differentiated cells were transferred to a 24 well-plate and ensured adherence. This transfer was necessary as cells grown on non-transparent MXene-coated surfaces were not visible under standard transmitted light microscopy. On day 7, the Micro group showed the highest osteogenic activity, with a normalized ALP activity of 2.62 relative to the uncoated planar control (*p* = 0.0023) (**Fig. 3b-c**). The coated planar group also showed notable osteogenic activity, with a relative ALP activity of 1.75, indicating the effectiveness of the MXene/HAP coatings in promoting initial cell adhesion and affecting early-stage cell differentiation. The other patterned groups, namely Nano and Submicro, displayed ALP activity levels similar to the uncoated planar control group. Extending the culture period to 14 days, the Micro group maintained its lead in osteogenic activity, with a relative ALP activity of 6.63 (normalized to the uncoated control at day 14; *p* < 0.0001). While the planar coated group showed an increase level of ALP activity compared to the planar uncoated control group, this was not statistically significant, a trend also observed in the Nano and Submicro groups (**Fig. 3c**). These findings suggest that microscale topographical cues have a more pronounced effect on osteogenic efficacy than chemical cues (i.e., HAP particles), and the osteogenic potential of topographical cues within the nano- to submicro-scale range is comparable to the chemical cues.

**Figure 3.**
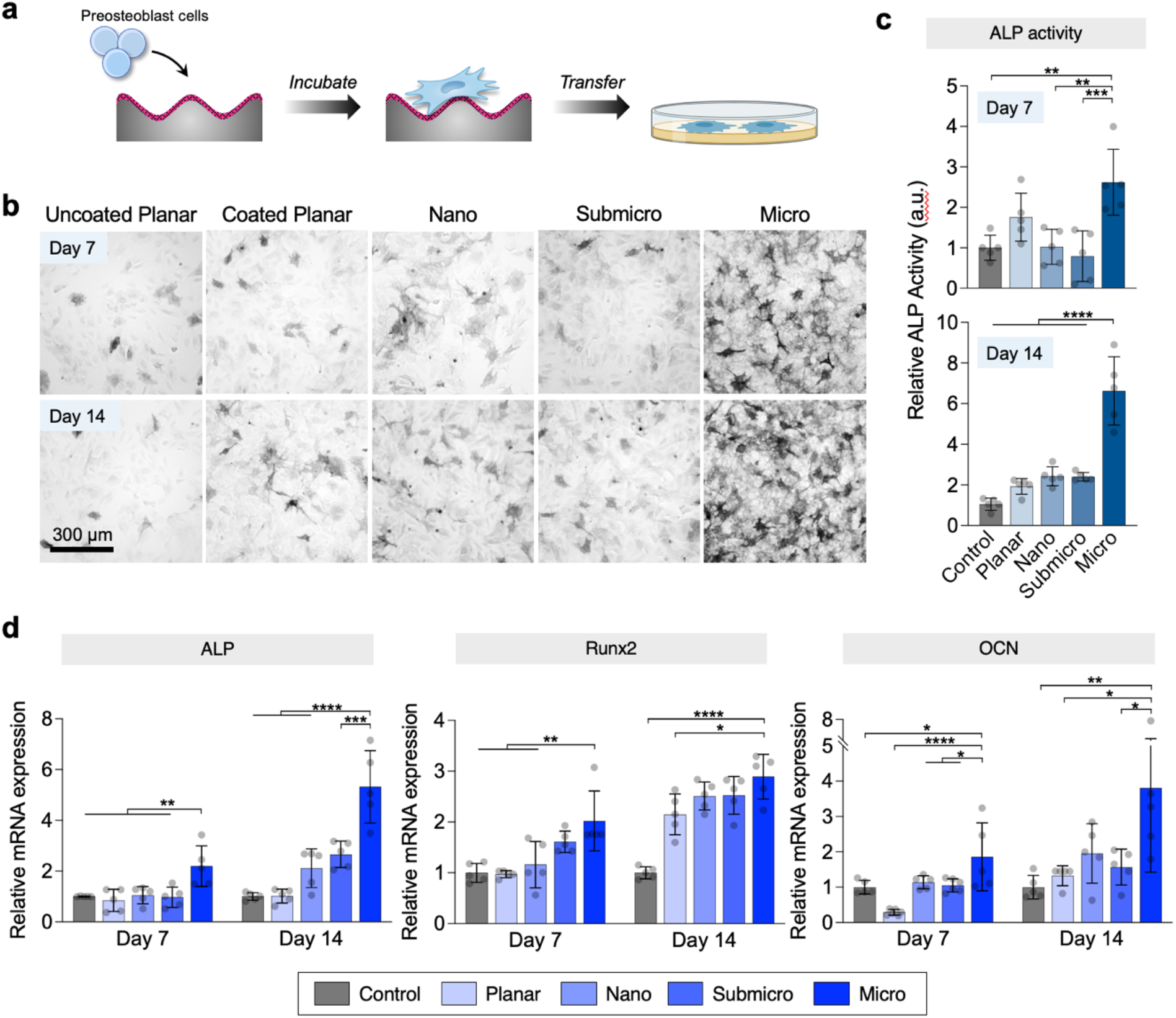
Comparison of osteogenic efficacy of flat controls and crumpled MXene multilayers. **(a)** Schematic of osteoblast culture setup for the evaluation of alkaline phosphatase (ALP) activity of the test samples. **(b)** Visual inspection of temporal expression patterns for ALP signals and **(c)** relative ALP activity of MC3T3-E1 cells cultured on different samples for 7 and 14 days. **(d)** Relative mRNA expression level of ALP, Runx2, and OCN of MC3T3-E1 cells cultured on different samples for 7 and 14 days. Each condition was tested with at least three biological replicates (error bar = standard deviations; *n* ≥ 5; ****p < 0.0001, \*\*\**p* < 0.001, \*\**p* < 0.01, and \**p* < 0.05), and at least three random images were taken from each sample.

To deepen our understanding of osteogenic differentiation at the gene level, we performed a quantitative polymerase chain reaction (qPCR) analysis focusing on three key osteogenic-related genes: ALP, runt-related transcription factor 2 (Runx2), and osteocalcin (OCN) after culturing cells for 7 and 14 days. A specific transcription factor to osteoblast differentiation, Runx2, can upregulate other osteogenic genes and initiate mineralization of immature osteoblasts.^46^ OCN is a marker of osteogenic differentiation later in the mineralization period.^47, 48^ It has pronounced involvement in controlling nuclear-encoded mitochondrial genes and developmental genes, contributing to the dynamic variations evident in transcriptional patterns during the differentiation process.^49^ At both days 7 and 14, micro scale topographies exhibited significantly elevated expression levels of ALP, Runx2, and OCN compared to the other groups (**Fig. 3d**). Specifically, on day 14, gene expressions in the Micro group were about 5.3 times higher for ALP, 3 times higher for Runx2, and 4 times higher for OCN, relative to the uncoated planar control. Overall, the findings from our cell differentiation studies strongly support our hypothesis that the MXene/HAP coated surfaces with micro-crumpled patterns significantly increase osteogenic differentiation of MC3T3-E1 cells. This effect is pronounced when compared to both uncoated and coated planar controls, highlighting the combined impact of chemical and topographical cues on osteogenesis.

### Macrophage response to different topographical cues

Osteoimmunomodulation plays a pivotal role in osteointegration by regulating immune cell responses and subsequent bone cell behavior.^50^ Upon implantation, the innate immune system reacts to the implanted materials, creating an immune niche that ultimately determines the fate of implant-to-bone integration.^51, 52^ As such, materials with immunomodulatory properties that promote an anti-inflammatory and prohealing immune microenvironment are crucial for enhancing osteointegration. Among the earliest immune cells to interact with implants are macrophages, which direct the host immune response. Remarkably, macrophages exhibit exceptional plasticity, adjusting their functions by polarizing into proinflammatory M1 or anti-inflammatory M2 phenotypes in response to local environmental cues.^53, 54^ Typically, M1 macrophages are associated with egg-shaped structures, while M2 macrophages often assume a more elongated form.^55, 56^ Given the established correlation between cell morphology and macrophage phenotype,^42, 57^ we reasoned that the micro-topographical cues are more effective at directing macrophages towards the M2 phenotype, due to enhanced cell adhesion and spreading, compared to the nano-topographical cues and flat controls. To test this hypothesis, we evaluated the morphology change and activation of macrophages on different samples.

To evaluate the immunomodulatory efficacy, Raw 264.7 macrophages were cultured on different test groups for 24 hours and then analyzed through (i) SEM for cell morphology, (ii) immunofluorescent staining assay for macrophage polarization, and (iii) gene expression analysis for macrophage activation and osteoinductive activity. As shown in **Fig. 4a-b**, macrophages in the Micro group exhibited more elongated morphologies and higher cell aspect ratios compared to other groups. To explore the influence of different test groups on macrophage polarization, we used immunofluorescent staining assay and measured the expression levels of CD206 and iNOS. These markers were chosen because iNOS (inducible nitric oxide synthase) is an important marker for M1 macrophage activation, while CD206 is indicative of M2 polarization.^58^ **Fig. 4c-d** show the results of immunofluorescent staining images and the quantitative fluorescence intensity of individual cells. For iNOS expression, all patterned groups (Micro, Submicro, and Nano) showed the lowest levels, ∼0.6 normalized to the uncoated planar control (*p* < 0.0001), with no significant variance among them. Conversely, for CD206 expression, the Micro group exhibited a relatively higher level than the other groups. These findings suggest that macrophages on topographically patterned surfaces are more likely to transition towards the M2 phenotype, while those on unpatterned surfaces predominantly shift towards the M1 phenotype.

**Figure 4.**
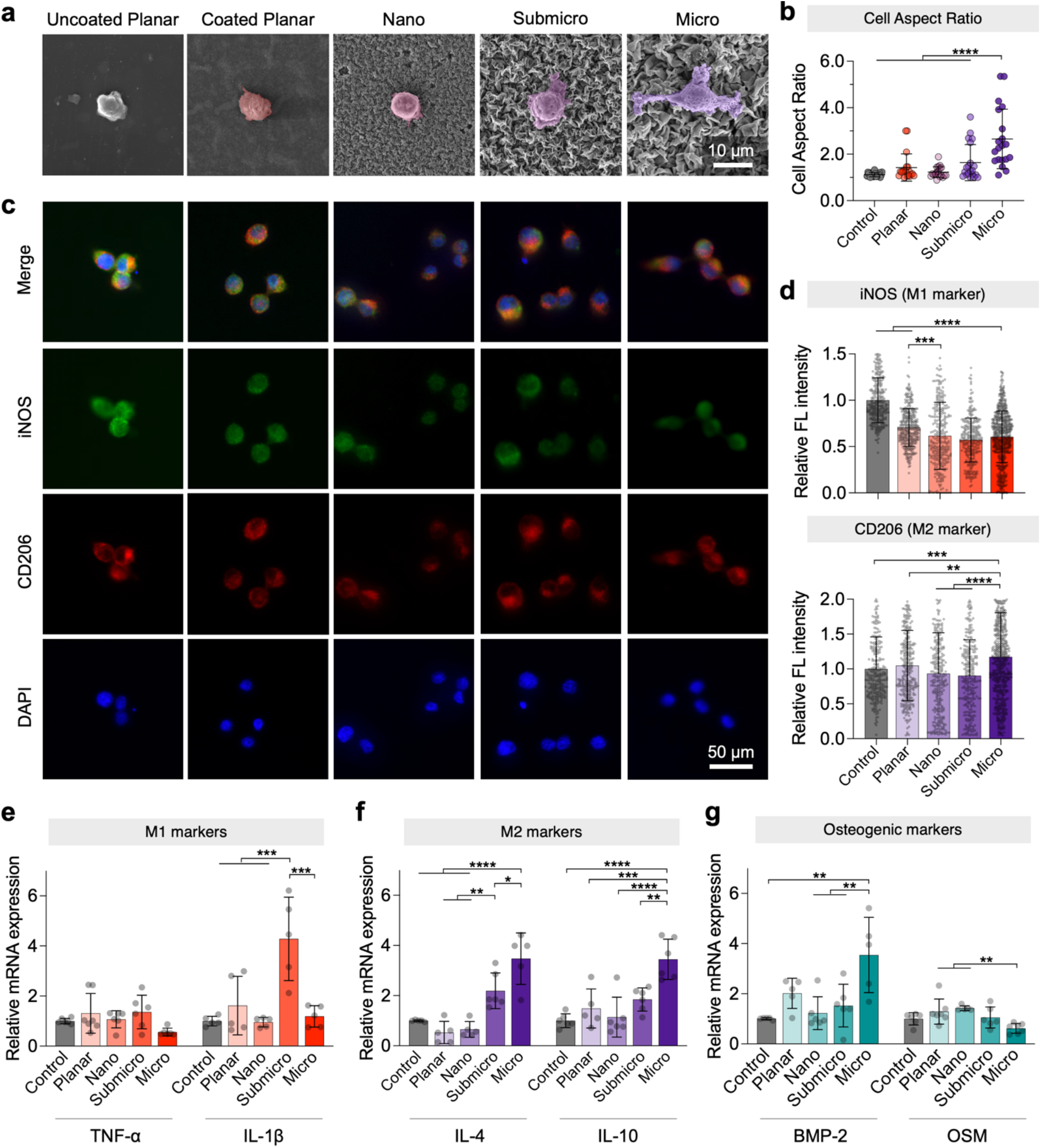
Comparison of immune responses of macrophages (RAW 264.7) on flat controls and crumpled MXene multilayers. **(a)** Cell morphologies of macrophages on different samples under SEM. **(b)** Relative cell aspect ratio of macrophages based on SEM images. **(c)** Representative fluorescence images of CD206 (red), iNOS (green), and nucleic (blue) staining of macrophages, and **(d)** Quantitative fluorescence intensities of CD206 and iNOS of macrophages on different samples. Relative mRNA expressions of **(e)** M1 macrophage-related genes (IL-1β and TNF-ɑ); **(f)** M2 macrophage-related genes (IL-4 and IL-10); and **(g)** osteogenic-related genes (BMP-2 and OSM). Each condition was tested with at least three biological replicates (error bar = standard deviations; *n* ≥ 5; \*\*\*\**p* < 0.0001, \*\*\**p* < 0.001, \*\**p* < 0.01, and \**p* < 0.05), and at least three random images were taken from each sample.

To further validate the phenotypes of polarized macrophages on different samples, we analyzed the expression levels of genes related to M1 and M2 macrophages (**Fig. 4f-g**). For M1-related gene TNF-ɑ, we found comparable expression levels across all groups with no significant differences. However, the expression of another M1-related gene, IL-1β, was significantly higher in the Submicro group than in other groups. In contrast, for M2-related genes (IL-4 and IL-10), there was a noticeable trend of increased expression correlating with the scale of crumpled topographies, from nano to microscale. Beyond their role in modulating inflammatory responses, macrophages also play an important role in bone regeneration by secreting osteoinductive factors, such as bone morphogenetic protein-2 (BMP-2) and oncostatin M (OSM). The expression levels of BMP-2 and OSM of macrophages cultured on different samples are shown in **Fig. 4h**. Consistent with previous reports, we observed that macrophages on the planar controls and those in the Nano group, which are M1 polarized, tend to secrete more OSM while a higher secretion of BMP-2 is associated with macrophages in the Micro group, which are M2 polarized.^59^ The trend of BMP-2 gene expression across the test samples paralleled that of M2-related genes (IL-4 and IL-10), while the OSM expression trend followed that of M1-related genes.

## DISCUSSION

In this study, we aimed to elucidate how controlled nano-/micro-topographies modulate cellular behaviors—including adhesion, spreading, morphology. Using our custom bottom-up topographic patterning method, we developed a series of surfaces with controlled topographical patterns ranging from nanometer to micrometer scales. These tailored surfaces served not only as distinct 2.5D physical topography cues but also as a robust platform for interrogating surface-cell interactions, important in both tissue growth and immunomodulatory contexts.

Our experimental design incorporated two distinct control groups: an uncoated planar and a MXene/HAP-coated planar surface. This approach enabled us to differentiate the cellular responses attributable to topographical and chemical cues. A key observation from our studies was the noticeable variance in cellular behaviors—namely cell adhesion, spreading, proliferation, and differentiation—across crumpled surfaces with different feature sizes. This highlighted the pivotal role of topographic pattern size in modulating cellular functions. Notably, the enhanced initial cell adhesion observed on all MXene multilayer coated surfaces, irrespective of the presence of crumpled structures, could be attributed to a synergistic effect of increased surface roughness and the osteoconductivity of embedded HAP particles.

We observed that cell morphology—a parameter including cell aspect ratio and spreading— emerges as a crucial determinant in the process of cell activation (e.g., osteogenic differentiation and macrophage polarization). This was particularly evident in the behaviors of MC3T3-E1 and RAW 264.7 cells, which exhibited markedly different spreading morphologies on the planar controls compared to the coated patterned surfaces. We observed filopodia on these surfaces formed initial attachments, evolving into lamellipodia and guiding cell spreading and morphology, in line with existing literature on pseudopodial dynamics.^60^

Furthermore, our study underscored that the micro-crumpled structures with a specific wavelength of ∼2 μm (comparable to the size of a lamellipodium) facilitated the formation of extended lamellipodia with minimal resistance to cell spreading. We found that cells cultured on micro-crumpled surfaces exhibited not only an elongated shape but also promoted proliferation and upregulated osteogenic differentiation, indicating that these changes were induced by topography cues. The early-stage osteogenic differentiation, as evidenced by increased ALP activity, was significantly more pronounced in cells cultured on the micro-crumpled surfaces. Our findings suggest a link between the abundance of filopodia/lamellipodia and osteogenic differentiation via cytoskeleton tension, likely through the RhoA-ROCK pathway.

Additionally, we explored the influence of surface topography on macrophage polarization. The microscale crumpled surfaces, offering abundant anchorage points, led to extensive filopodia formation and spreading. This response to topographical cues likely activated the Rho family of GTPases, subsequently triggering the RhoA/ROCK signaling pathway, thereby inducing M2 macrophage activation.^61, 62^ We postulate that microscale topographies can shift macrophages towards M2 polarization, creating a favorable immune microenvironment for osteogenic activity.

In summary, our study elucidates the profound influence of MXene/HAP-coated surfaces with controlled topographical patterns on regulating osteogenic differentiation in preosteoblast cells as well as macrophage polarization. While further investigation is needed to fully understand the underlying mechanisms and the involved signaling pathways and to assess in vivo efficacy, our findings establish the promise of topographically patterned MXene/HAP multilayers as an efficient platform with both osteogenic and immunomodulatory properties that can be fabricated in a low-cost, scalable manner and be applicable for a range of biomaterials and implantable devices. Our results not only validate the potential of these coatings but also provide essential insights that could inform the development of next-generation biomaterials, specifically tailored for enhanced tissue regeneration and immune responses.

## METHODS AND EXPERIMENTAL

### Materials and cell lines

Ti_3_AlC_2_ MAX phase powders (300 mesh, ≥99%) were purchased from Laizhou Kai Kai Ceramic Materials Co. Clear heat-shrinkable polystyrene (PS) films were purchased from Grafix. Ultra-pure deionized water was used (18.2 megohm, Mill-Q pore). All other materials were sourced from Sigma Aldrich or Thermo Fisher Scientific unless otherwise noted. All reagents were used as received, without further purification.

Cell lines used in this study include MC3T3-E1 pre-osteoblast cells and RAW 264.7 macrophages. All cell lines were purchased from ATCC. MC3T3-E1 was maintained in Minimum Essential Medium α (αMEM), and RAW 264.7 was maintained in Dulbecco’s Modified Eagle’s Medium (DMEM). All media were supplemented with 10% fetal bovine serum and 1% penicillin-streptomycin (Corning). Cells were cultured in a humidified chamber at 37C with 5% CO_2_, and subcultured at ∼80% confluence using 0.05% Trypsin-EDTA. All cell lines were routinely tested using MycoAlert mycoplasma detection kit (Lonza).

### Synthesis of Ti_3_C_2_T_x_ MXene

Ti_3_C_2_T_x_ MXene nanosheets were synthesized using a modified version of a previously reported method involving in situ hydrofluoric acid etching (**Fig. S1**). To initiate the synthesis, Ti_3_AlC_2_ powder (1.0 g), a type of Max phase material, was subjected to etching using a solution comprising LiF (7.5 g) and HCl (9 M, 40 mL). The etching process was carried out under stirring for 24 hours to remove the aluminum layers from Ti_3_AlC_2_. After the etching process, the resulting residue was obtained through centrifugation and subsequently washed multiple times with deionized (DI) water until the pH value reached approximately 6.0. After that, the residue was mixed with 100 mL of DI water and subjected to bath sonication for 1 hour at 4 ºC. After sonication, the mixture was centrifuged at 3500rpm for 30 minutes to separate the heavier components. The supernatant obtained from this process contained the stable Ti_3_C_2_T_x_ MXene nanosheets.

### Fabrication of crumpled MXene multilayers

First, MXene/HAP multilayers were deposited on biaxially-oriented polystyrene (PS) films. Briefly, HAP nanoparticles suspension (10 wt%) was diluted 5 times in ultra-pure milliQ water. The resulting mixture was then passed through a sterile 0.45 μm cellulose membrane for filtration. The MXene solution was diluted to 1 mg/mL in water. All substrates were cleaned by sequential sonication in ethanol and water for 15 min each. After washing, the substrate was dried with nitrogen and treated with oxygen plasma (Tergeo-Plus, PIE Scientific) for 10 min at 50 W under vacuum. For the dip-LbL based multilayer assembly, we used an automated dipping robot (KSV NIMA Dip coater, Biolin Scientific) and the Ti_3_C_2_T_x_ MXene and HAP solutions. The plasma-treated substrates were first immersed in PDAC solution for 30 min to deposit a base layer, and rinsed with DI water twice for 2 min each to remove the weakly bound molecules. After that, bilayer multilayers were fabricated by alternate dipping in HAP solution for 5 min followed by two consecutive rinse steps in DI water baths twice for 2 min each, and then into MXene solution for 5 min followed by the same rinse cycle. This cycle made one bilayer of (MXene/HAP)_1_, and the bilayer was repeated to fabricate the desired multilayers, denoted as (MXene/ HAP)_*n*_ where *n* is the bilayer number. After LbL assembly, mechanical nanomanufacturing was used to create crumpled topographical patterns. The prepared MXene/HAP multilayers on PS substrates were cut into a desired size and shrunk by heating in an oven at 130 °C for 5 min. The strain of the PS film was determined using the equation ε = (Li – Lf)/Li. The as-prepared and crumpled multilayer films were stored in a closed container prior to subsequent analysis.

### Materials characterization

Film thickness and surface roughness were determined by AFM (Dimension Icon, Bruker Corp.) in tapping mode as well as UV-vis spectrophotometer (Agilent Cary 8454). For AFM measurement, MXene multilayers grown on glass were scratched with a plastic razor blade, and thickness was measured at three predetermined locations. Surface morphology of the crumpled surfaces was observed using SEM (Tescan Mira3).

### Immunostaining assay for preosteoblast cells

For cell proliferation testing, MC3T3-E1 cells were cultured on test samples—including the uncoated- and coated planar films, nano, submicro, and micro-scale topographies—with a cell density of 1×10^4^ for 1, 3, and 5 days. After the respective incubation periods, the cells were fixed with 4% paraformaldehyde for 15 min and washed with PBS. For cell proliferation testing, MC3T3-E1 cells were seeded onto the test samples at a cell density of 2×10^4^ and incubated for 1 day. Following this, cells were fixed with 4% paraformaldehyde and then permeabilized with 0.1% Triton X-100 in PBS for 15 min, followed by washing with PBS. Subsequently, cells were incubated with Alexa Fluor 488 Phalloidin, along with 1% bovine serum albumin (BSA), for 30 min at room temperature in the dark.

### Immunostaining assay for macrophages

RAW 264.7 cells were seeded on the test samples at a density of 10^4^ cells and incubated for 24 hours. Following this, cells were fixed and treated in a permeabilization/blocking buffer (PBS supplemented with 5% goat normal serum and 0.3% Triton X-100) for 60 min. After removing the perm/blocking buffer, a cocktail of CD206 monoclonal antibody (MR5D3) rat/ IgG2a (1:200) and iNOS rabbit IgG monoclonal antibody (1:200) in a staining buffer (PBS supplemented with 1% BSA and 0.3% Triton X-100) was added and incubated overnight at 4 ºC in the dark. The next day, cells were washed with PBS and stained with secondary antibodies—Anti-rabbit IgG Alexa Flour 488 (1:500) and Anti-rat IgG Alexa Flour 647 (1:500) —at room temperature for 1 hour.

### Quantitative polymerase chain reaction (qPCR)

Total RNA from cells was extracted using TRIzol (Invitrogen) and then reverse-transcribed into cDNA utilizing a PrimeScript RT Master Mix (Takara Bio, Japan), following the manufacturer’s guideline. Real-time PCR was performed on a Quant Studio 3 RT-PCR system for internal reference for glyceraldehyde-3-phosphate dehydrogenase (GAPDH); (i) osteogenic factors Runx2, ALP and OCN for preosteoblast cells; and (ii) M1 markers IL-1β and TNF-α, M2 markers IL-4 and IL-10, and immune osteogenesis-related marker BMP-2 and OSM. The primers used are summarized in **Table S2**.

### Alkaline Phosphatase (ALP) assay

To assess the osteogenic efficacy of test samples, we performed ALP activity assay. For each group, MC3T3-E1 cells were seeded at a density of 2 × 10^4^ and incubated for 7 and 14 days. Osteogenic induction medium consisting of growth medium with 50 μM L-ascorbic acid, 10 mM - glycerol phosphate and 50 nM dexamethasone was used. After 7 and 14 days, cell were washed with PBS, trypsinized, and transferred to a 24 well-plate. The transferred cells were incubated for a few hours to adhere and then fixed. ALP activity then was measured using an Alkaline Phosphatase Colorimetric Assay kit (Abcam), which quantifies the ALP enzyme activity, based on the manufacturer’s specifications.

### Cell morphological observation via SEM

Cells were seeded on the films (uncoated- and coated-films, nano, submicro, and micro-scale topographies) at a desired density of 1-2 × 10^4^ for 24 hours. The samples were then washed with PBS, and fixed with 4% paraformaldehyde for 15 min. After washing the samples with PBS, they were incubated with 1% osmium tetroxide in PBS for 30 min. The fixed samples were dehydrated using 30%, 50%, 70%, 100%, 100% with each step lasting 5 min and then dried using a critical point dryer (Leica EM CPD300). The samples were sputter coated with gold, and then imaged using the Tescan Mira3 SEM.

### Microscopy and image analysis

Fluorescence images of immunostained cells were acquired on Axio Observer fluorescence microscope (Carl Zeiss Inc.). All cells were stained with DAPI and visualized after capture with a DAPI filter cube. Data were analyzed using ImageJ and Cellprofiler.

### Statistics

Statistical analyses and data plotting were performed in GraphPad Prism 10. For correlations, the linear least squares fitting was performed at the 95% confidence level, and the Pearson correlation coefficient (*r*) was used to quantify the correlations between different variables. Group differences were tested using the nonparametric t-test for two groups and analysis of variance (ANOVA) with post hoc analysis for more than two groups. All tests were two-sided, and *p* < 0.05 was considered statistically significant.

## Data availability

The data that support the findings of this study are available from the corresponding author upon reasonable request.

## Supplementary Materials

Table S1: Comparison of osteogenic and immunomodulatory activities of various topographically patterned surfaces

Table S2: Primer sequences for qPCR analysis.

Fig. S1: MXene synthesis and characterization.

Fig. S2. Self-assembly formation of MXene/HAP hybrids.

Fig. S3. Growth curve of (MXene/HAP)_*n*_ films.

Fig. S4. Morphological characterization of (MXene/HAP) multilayers with crumpled structure.

Fig. S5. Mechanical stability testing.

Fig. S6. Film degradation testing.

## Acknowledgements

This work was supported by a Startup Grant from University of Michigan, College of Engineering. We would like to thank Haiping Sun for assistance with SEM and AFM.

## Author contribution

M.A. and J.M., conceived the study. M.A. performed experiments. M.A. and J.M. analyzed data. M.A. and J.M. wrote and edited the manuscript.

