## Supplementary materials for "Designer micro/nano-crumpled MXene multilayer coatings accelerate osteogenesis and regulate macrophage polarization"

**Table S1**. Comparison of osteogenic and immunomodulatory activities of various topographically patterned surfaces.

| Material | Pattern | Dimensions (height; diameter; density; period) | Patterning techniques | Precision | Scale-up capability | Osteogenic activity | Immuno modulatory activity | Ref |
|---|---|---|---|---|---|---|---|---|
| **$TiO_2$** | Honeycomb | 0 nm; 90-5000 nm; n/a; n/a | 3D imprinting | High (nm) | Low-Med | Nano | n/a | 1 |
| **$TiO_2$** | Honeycomb | 0 nm; 90-5000 nm; n/a; n/a | 3D imprinting | High (nm) | Low-Med | Nano | Nano | 2 |
| **Zinc plates** | Grooves | 80-1200 nm; n/a; n/a; n/a | Chemical etching | Low (µm-mm) | High | Micro | Micro | 3 |
| **$TiO_2$** | Hierarchy | 400- 1500 nm; 0-500 nm; n/a; 500- 1500 nm | Acid etching | Low (µm-mm) | High | Micro | n/a | 4 |
| **IP-L780 resin, glass** | Pillars | 250-1000 nm; 250 nm; 100-205 nm; 700-1000 nm) | Laser printing | High (nm) | Low | Micro | n/a | 5 |
| **$SiO_2$, $TiO_2$, $CrO_3$, $Al_2O_3$** | Wrinkles | 250-2500 nm; n/a; n/a; 1-80 µm | Pre-stretching, thermal evaporation | Med (nm-µm) | Med | Micro | n/a | 6 |
| **Ti** | Grooves | 800-1300 nm, n/a, n/a, 200-50000 nm | Lithography | High (nm) | Low | n/a | Micro | 7 |
| **PDMS** | Wrinkles | 50-4300 nm, n/a, n/a, 500 nm-27 µm | Imprinting | High (nm) | Low-Med | Micro | n/a | 8 |
| **PLA, PCL, PDMS** | Gratings | 114-1972 nm; n/a; n/a; 250 nm-2 µm | Lithography | High (nm) | Low | n/a | No pattern | 9 |
| **MXene, HAP** | Crumpled | 200-1000 nm; n/a; n/a; 300-2000 nm | Bottom-up manufacturing | High (nm) | High | Micro | Micro | This work |

**Table S2**. Primer sequences for qPCR analysis.

| Target | Forward primer | Reverse primer |
|---|---|---|
| ALP | GGACCATTCCCACGTCTTCAC | CCTTGTAGCCAGGCCCATTG |
| RUNX2 | CGCCTCACAAACAACCACAG | CGCCTCACAAACAACCACAG |
| OCN | AGCAGTTGGCCCAGACCTA | TAGCGCCGGAGTCTGTTCACTAC |
| TNF-$\alpha$ | CCCTCACACTCAGATCATCTTCT | GCTACGACGTGGGCTACAG |
| IL-1$\beta$ | TGCCACCTTTTGACAGTGATG | AAGGTCCACGGGAAAGACAC |
| OSM | CCCGGCACAATATCCTCGG | TCTGGTGTTGTAGTGGACCGT |
| IL-10 | GCTCTTACTGACTGGCATGAG | CGCAGCTCTAGGAGCATGTG |
| IL-4 | GGTCTCAACCCCCAGCTAGT | GCCGATGATCTCTCTCAAGTGAT |
| BMP-2 | TGAGGATTAGCAGGTCTTTGC | GCTGTTTGTGTTTGGCTTGA |
| GAPDH | AGGTCGGTGTGAACGGATTTG | TGTAGACCATGTAGTTGAGGTCA |

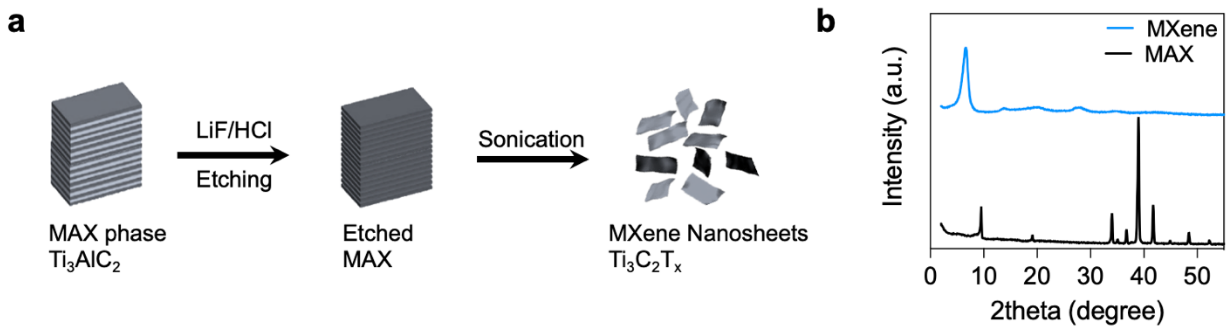


**Figure S1. MXene synthesis and characterization.** (a) Schematic illustration of $Ti_3C_2T_x$ MXene synthesis: Selective etching of Al atoms from $Ti_3AlC_2$ MAX phase using LiF and HCl mixture. (b) XRD patterns of powder $Ti_3AlC_2$ MAX and $Ti_3C_2T_x$ MXene: multilayer and nanosheet papers were fabricated through vacuum filtration of the MXene solution. Following exfoliation and delamination, the distinctive (002) peak of $Ti_3AlC_2$ shifted from 9.5º to 6.8º in the MXene nanosheets.

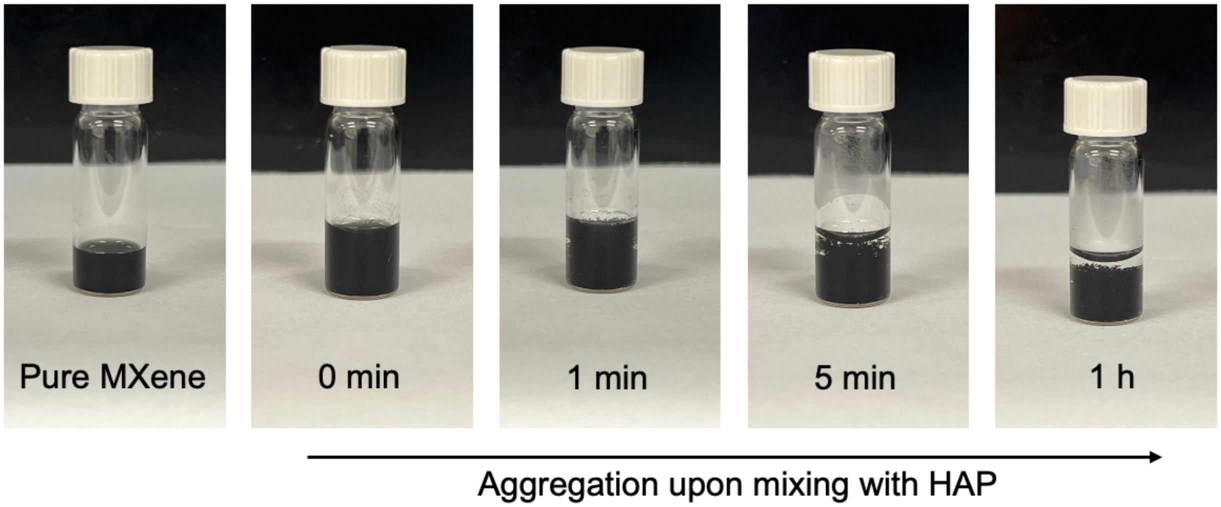


**Figure S2**. **Self-assembly formation of MXene/HAP hybrids**, resulting from the mixture of MXene dispersion (1 mg/mL)_with HAP nanoparticle solution (2 wt% of 112 nm HAP) at a 1:1 volume ratio. Digital photos show the hybrid formation at various timepoints after mixing, arranged from left to right.

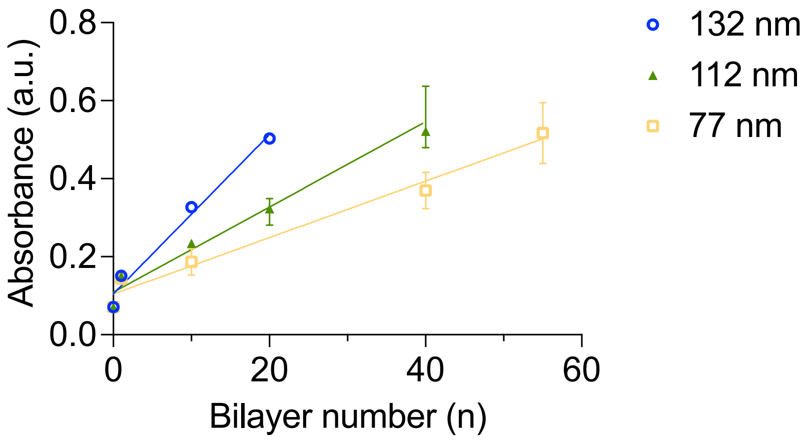


**Figure S3**. **Growth curve of (MXene/HAP)$_n$ films** as a function of bilayer number (*n*), measured by UV-vis spectroscopy. The rate of film growth is dependent on the size of HAP particles, with increasing trend observed as the size of HAP ranges from 77 nm to 132 nm.

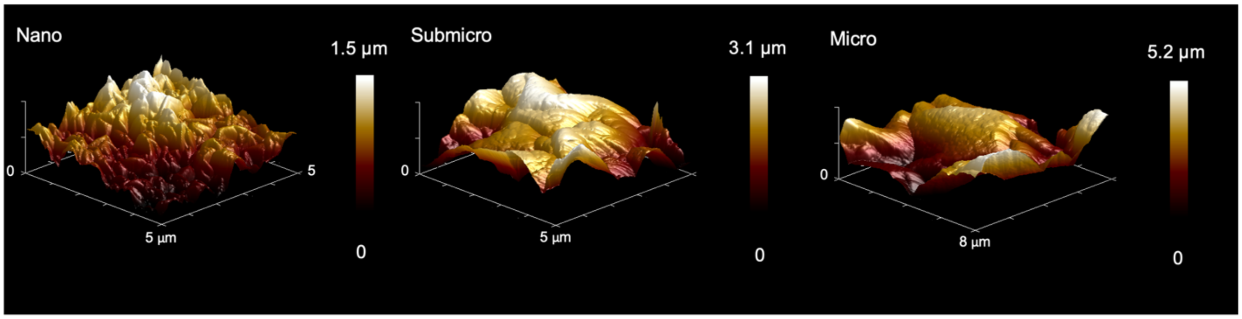


**Figure S4**. **Morphological characterization of crumpled MXene/HAP multilayers**. 3D height AFM images show the crumpled structures of MXene/HAP multilayer coatings on PS substrates, featuring nano-, submicro- and micro-structures.

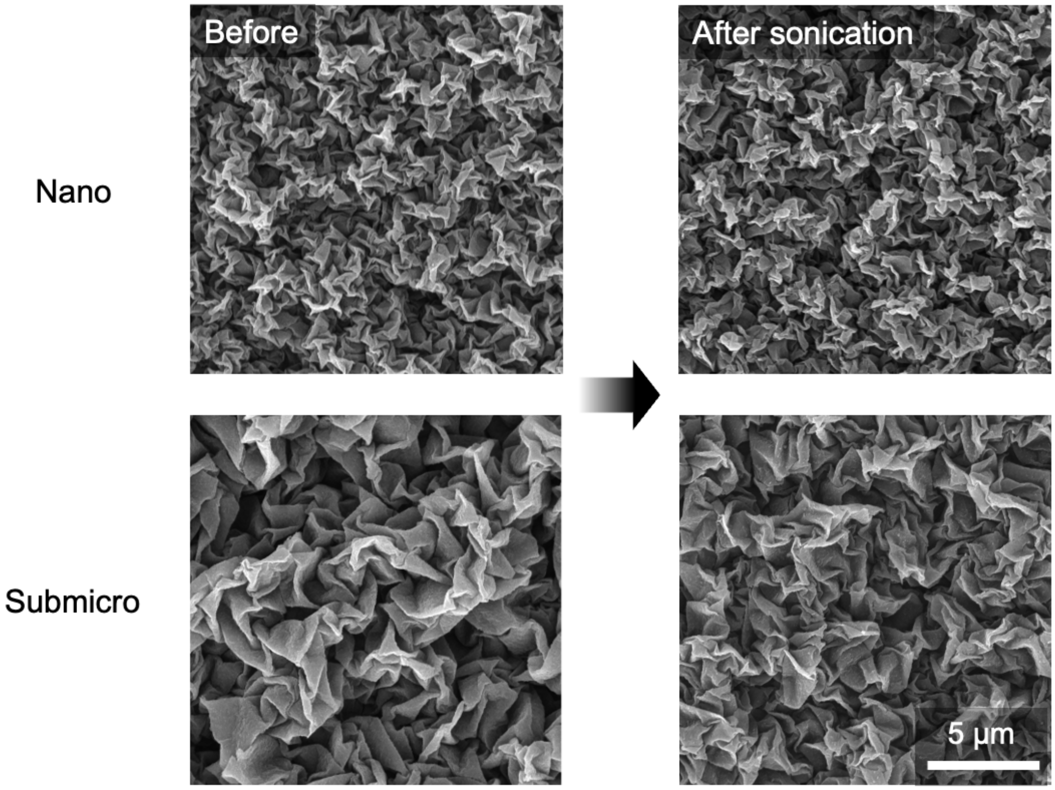


**Figure S5**. **Mechanical stability testing**. The nano- and submicro-patterned surfaces were subjected to a 15-minute batch sonication in water, with no obvious morphological distortion observed.

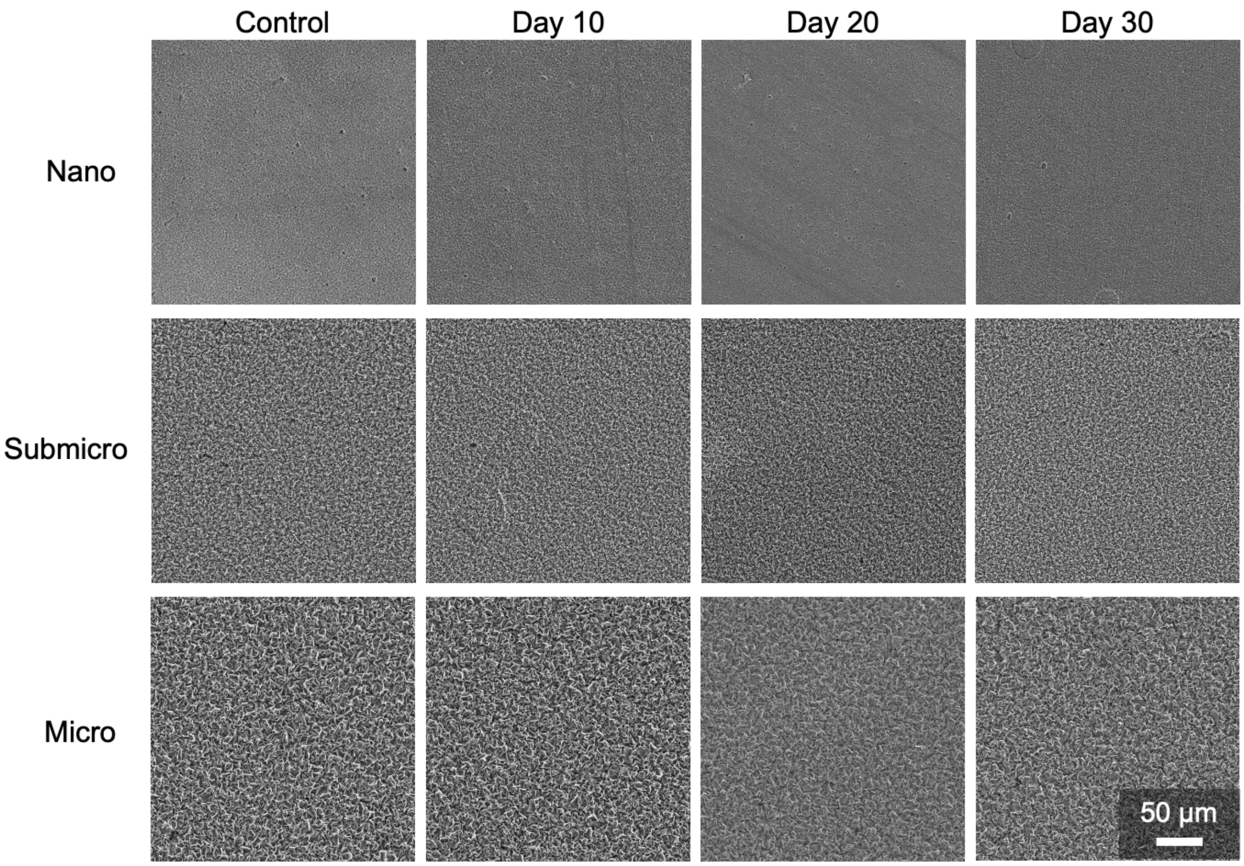


**Figure S6**. **Film degradation testing.** The topographically patterned surfaces were incubated in cell culture media at room temperature, and no obvious morphological distortion was observed over a period of 30 days.
